# Development and Characterization of a FRET-based Formin Tension Sensor in Living Cells

**DOI:** 10.64898/2026.07.11.737992

**Authors:** Philip Bleicher, John A. Hammer, James R. Sellers, Anjelika Gasilina

**Affiliations:** Laboratory of Molecular Physiology, Cell and Developmental Biology Center, National Heart, Lung and Blood Institute, National Institutes of health, Bethesda, MD, 20814, USA; Molecular Cell Biology Laboratory, Cell and Developmental Biology Center, National Heart, Lung and Blood Institute, National Institutes of health, Bethesda, MD, 20814, USA

## Abstract

Mechanotransduction *via* the actin cytoskeleton is linked to fundamental cellular processes such as morphogenesis, cell division, and motility, requiring the control of tensile forces mediated by the motor protein non-muscle myosin 2 (NM2). Formins such as mDia1 have been shown to elongate actin structures that are under mechanical tension; conversely, mDia1’s elongation rates are modulated by the applied force. Despite their relevance at the membrane/cortex interface, reported values for tension in formin-elongated actin filaments stem from theoretical estimates and simulations, but have not been amenable experimentally so far. Thus, we developed a Förster resonance energy transfer (FRET)-based, tension-sensitive probe (mDia1TS) and quantified the measured tension in live U2OS cells using fluorescence lifetime imaging microscopy (FLIM). Through whole-cell ROI analysis we show a short and long lifetime component, reporting an intensity-weighted, averaged lifetime corresponding to ∼3.5 pN. Upon mitogen stimulation of cells using EGF, we show that the tension homeostasis changed significantly, with a measurable increase in tension in the cell’s periphery and relaxation in its center. Furthermore, the reported average tension relaxed by 2 pN after adding the NM2 inhibitor para-nitroblebbistatin. We utilized siRNA knockdowns of individual NM2 paralogs (NM2-A, NM2-B, or NM2-C) to measure their individual contribution, revealing NM2-A as the main paralog to produce tensile force in this system. Taken together, we demonstrate that mDia1TS is able to directly determine that active mDia1 in cells is under tension, and that subcellular quantification with pN precision is possible.

**Significance:** Despite the fundamental importance of formins in regulating actin-based processes, reported values for tension in formin-mediated actin structures stem from simulations and theoretical estimates. In this study we developed a FRET-based, tension-sensitive reporter probe for formin mDia1, which we termed mDia1TS. Given the expanding clinical spectrum of DIAPH1/mDia1 mutations, our tool mDia1TS provides a quantitative tool for elucidation of changes in cytoskeletal assemblies.

## Introduction

The actin cortex is a contractile meshwork beneath the cell membrane, consisting of actin filaments, myosins and binding proteins^1-3^. To meet local, biomechanical demands, actin is dynamically remodeled, supporting cell shape, adhesion, migration, and cytokinesis^4^. Among cortical nucleators of actin, formins elongate unbranched filaments by processively recruiting monomers to growing barbed ends^5^. MDia1 is a key, ubiquitously expressed formin, known for its roles in cytokinesis, endocytosis, stress fiber and filopodia formation. In the cytosol, the mDia1 dimer adapts an autoinhibited state *via* interaction of its autoregulatory domains; however, RhoA-GTP binding and membrane interactions relieve this autoinhibition^6-8^.

In the activated state, the actin binding domains, named formin-homology domain 1 (FH1) and formin homology domain 2 (FH2), cooperate to mediate processive actin elongation^9-12^. The FH2 domains hereby form a head-to-tail dimer, encircling the barbed end of the growing filament. Some studies suggest that elongation occurs via a “stair-stepping” mechanism, where the two FH2 domains alternate between an open and closed state^13-15^. Addition of monomers in the closed state is restricted, with the FH2 domain actin as a “cap”, resulting in a dynamic gating mechanism^16^. The FH1 domain on the other hand has multiple binding sites for complexes of monomeric actin and profilin; and is largely unstructured^17^. Hence, it recruits actin beyond the diffusion limit, enabling the rapid elongation rates, for which mDia1 is known^18,19^.

*In vitro* studies have shown that actin elongation by mDia1 is force-sensitive, showing increased elongation rates under higher tension, within a range of few piconewtons of load^13,20,21^. This is explained by the FH2 domain’s gating mechanism, as tension likely stabilizes the open confirmation, which facilitates the incorporation of monomers^22^. It is also possible that load reduces the likelihood of backwards stepping. *In vivo*, direct study of tension and elongation rates is more challenging. However, mDia1 polymerizes actin in stress fibers, lamellipodia and filopodia, all of which are structures known to be under mechanical load^5^. A key regulator of tension in cells is non-muscle myosin 2 (NM2) and subsequently, its activity has shown to have a direct effect on elongation dynamics^23-26^. Although this interaction is known, is has not been possible to directly demonstrate that mDia1 is under load *in cellulo*, due to experimental limitations. Consequently, most numbers for tensile force used in mechanical models of the actin cortex stem from theoretical estimates^27,28^ or measurements of bulk forces, such as micropipette aspiration, atomic force microscopy (AFM) or optical tweezers, ranging from 15-1600 pN/µm^29^.

Here we developed an mDia1 tension sensor for live cells based on Förster resonance energy transfer (FRET). A FRET pair connected by a flexible linker^30,31^ is inserted directly before the unstructured FH1 domains of mDia1. If mDia1 is under load in cells and the FH1 domains are extended, this should stretch the linker and increase the distance between the FRET pair, leading to a measurable decrease in the FRET efficiency compared to unloaded conditions. Due to the ubiquitous nature of mDia1, this could address various open questions, such as how force fluctuates at the intersection of membrane and cortical actin during migration^32^. Another application are force measurements during apical constriction in the context of endocytosis, where mDia1 is known to be involved^33,34^. Furthermore, mDia1 localizes to filopodia^5^, which are actively remodeled *via* both actin polymerization and tensile force mediated by myosins, which could be quantified using this probe.

Using the mDia1 tension sensor (mDia1TS), we directly demonstrate that formin is under load in cells; and we quantify this load with pN precision. The average tension throughout the cell was 3.5 pN. We added para-nitroblebbistatin, an inhibitor of NM2 activity, which reduced the tension to 1.5 pN, but not to zero. Then, we knocked down the three paralogs of NM2, named NM2-A, NM2-B and NM2-C, individually. The NM2-A knockdown showed the strongest effect, reducing the average tension to 2.2 pN, while NM2-B and NM2-C reduced the tension to 2.7 pN and 2.6 pN respectively. Additionally, we showed that activating cells using epidermal growth factor (EGF) led to a measurable remodeling of tensile forces throughout the cell. Image fits of micropatterned cells show that stimulated cells display increased tension in their periphery, while simultaneously relaxing towards the center. Interestingly, the average tension throughout the cell as a whole was not affected by mitogen stimulation.

## Results

### Design of the mDia1 tension sensor

In cells, mDia1 forms a dimer that is activated *via* binding of RhoA·GTP to its N-terminal GTPase-Binding Domain (GBD) (Fig. 1A). N-terminal interactions also mediate the membrane localization, thus preventing the release of mDia1’s autoinhibition while soluble in the cytosol. The Diaphanous Inhibitory Domain (DID) regulates this process by interacting with the C-terminal Diaphanous Autoregulatory Domain (DAD), blocking the actin barbed end binding regions. The region after the DID, including the Coiled-Coil (CC) region mediate dimerization^5,11^. MDia1TS’s FRET pairs, consisting of a teal fluorescent protein TFP donor and a Venus acceptor, are located directly after these N-terminal domains. The FRET pair is connected by a flexible linker, leading to a high FRET state when no tension is applied and the linker is relaxed. Adjacent to the FRET pair is the FH1 domain, containing binding sites for profilin bound actin monomer. The two FH2 domains form a ring-shaped structure that wraps around the barbed end of the bound actin filament. When tension is applied to the actin filament, this force should be transmitted via the FH1 domains and lead to a measurable decrease in FRET efficiency as the linker is tensed (Fig. 1B). Hence, FRET% reported by mDia1TS quantifies the tension via actin towards the membrane at this critical intersection.

**Figure 1.**
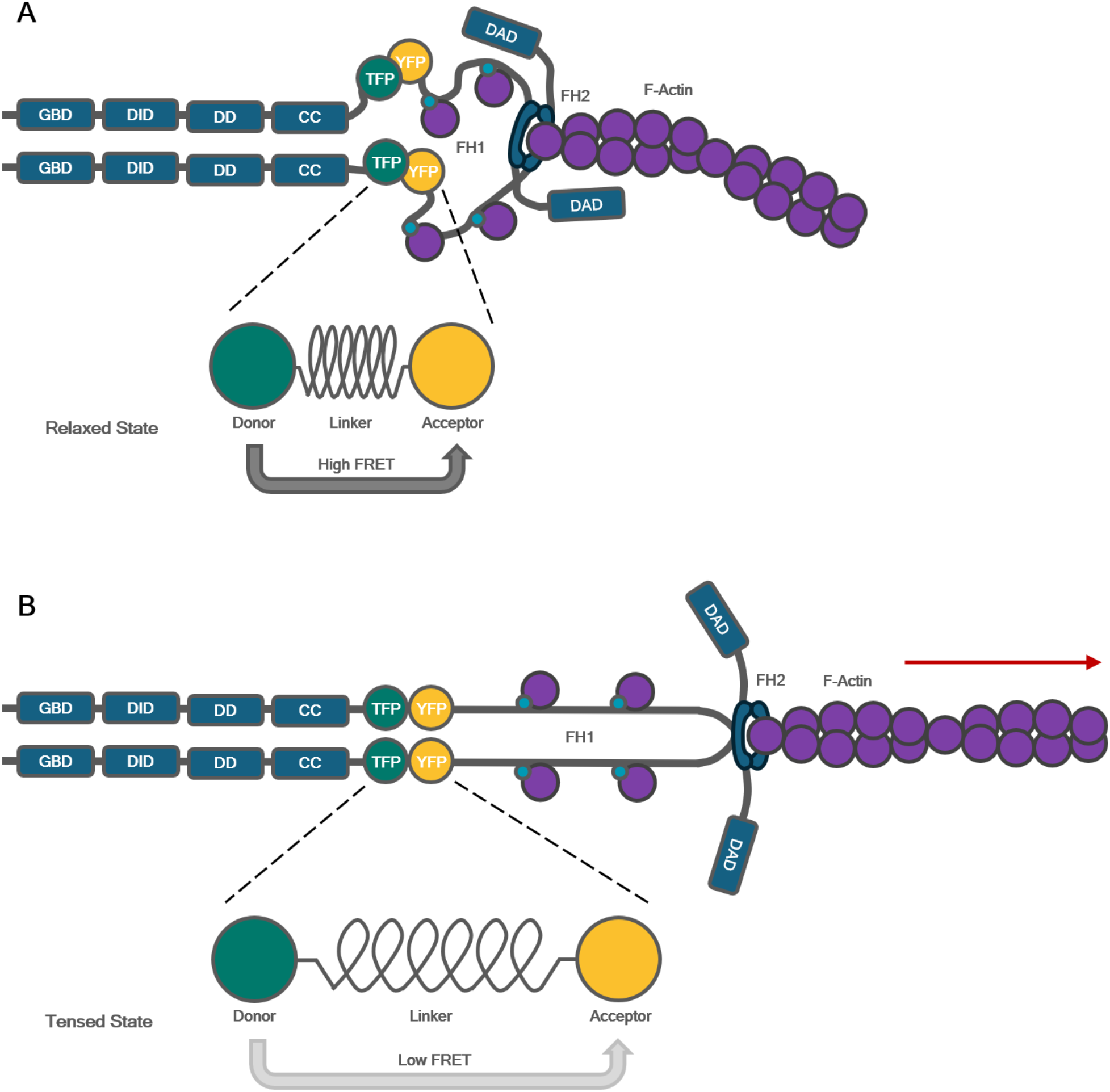
Design of the mDia1 tension sensor. The tension sensor module (TSMod) consists of a FRET donor (TFP) and acceptor (YFP) connected by a flexible linker. TSMod is located between the N-terminal domains (GBD, DID, DD, CC) and the C-terminal domains (FH1, FH2, DAD). Actin is shown in magenta; attached blue circles denote profilin-bound monomers. In the relaxed state, the FRET pair is in close proximity reporting relatively high FRET%. **A –** Schematic of the tension sensor module in its relaxed state. **B –** schematic of the tension sensor module in its tensed state. Tensile force mediated via bound, growing actin filaments extend TSMod, leading to a decrease in FRET%.

### Pharmacological inhibition of myosin motor activity or microtubule polymerization increases FRET efficiency of wild type full-length mDia1aTS

We used U2OS cells as it is a well characterized and established model for studying actin assembly, mechanotransduction and force generation^35,36^. We examined the cells using a confocal microscope with a FLIM module, to quantify both mDia1TS’s donor intensity to evaluate the sensors expression levels; and donor lifetime, which was used to calculate FRET%. For all images, the focus was set to the fluorescent layer of mDia1TS closest to the coverslip glass to minimize photons from unbound, cytosolic mDia1TS that is likely in an autoinhibited and thus relaxed state. Raw fluorescence images showed that mDia1TS expressed ubiquitously throughout the cell, with generally lower levels under the nucleus and at the cell’s periphery. Differences in lifetime levels were more subtle but showed a trend of elevated lifetimes near the periphery and under the nucleus. Using cytoskeletal drugs, lifetime levels varied much clearer compared to untreated cells, with a notable decrease in cells treated with para-nitroblebbistatin (pnBB) or the microtubule polymerization inhibitor nocodazole, and a lifetime increase in cells treated with CK666, an inhibitor of the Arp2/3 complex (Fig. 2A). These lifetime image fits showed that subcellular localization of varying tension levels from single cells are possible, but not easily amenable due to the signal-to-ratio and relatively subtle differences throughout single cells. Thus, lifetimes were fitted using whole-cell ROIs and comparing populations of at least 40 cells.

**Figure 2.**
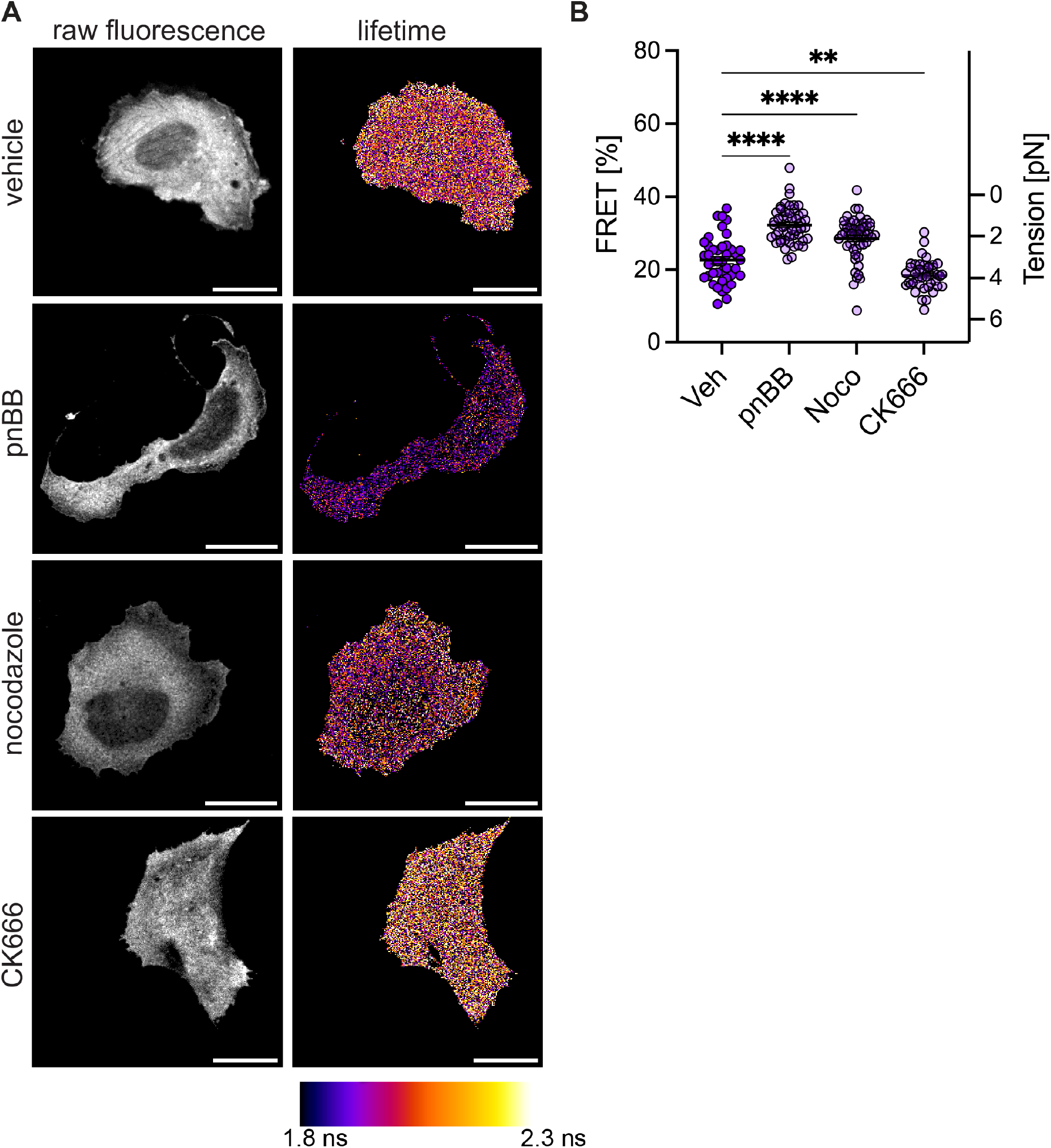
Pharmacological inhibition of myosin motor activity and microtubule polymerization increases FRET efficiency of wild type full-length mDia1TS. **A –** Representative raw fluorescence (left) and lifetime color-coded (right) images of U2OS cells expressing wild type full-length mDia1TS treated with vehicle control, para-nitroblebbistatin (pnBB), nocodazole, or CK666. **B –** Graph summarizing mean ± SEM of FRET percentage (% FRET) for each treatment condition. Each data point represents the FRET% obtained from a fit from a single, whole-cell ROI. Tension was estimated using a published calibration curve^31^. N = 50 – 60 cells per treatment, **** p <0.0001 based on ordinary one-way ANOVA with Dunnett’s multiple comparisons test. Scale bar = 20 µm.

In para-nitroblebbistatin treated cells (pnBB), the FRET efficiency increased to 32.26 ± 0.71% compared to vehicle (Veh) treated cells, which exhibited FRET efficiency of 22.60 ± 0.97%. Treatment with nocodazole, likewise increased the FRET efficiency to 28.56 ± 0.85%. Treatment with CK666 decreased FRET efficiency to 18.43 ± 0.66% (Fig. 2B).

Tension levels were calculated using a published calibration of the tension sensor module and using a correction established for the specific donor/acceptor FRET pair TFP/Venus (Grashoff, Hoffmann, Bleicher). We estimate the tension of Veh treated cells at 3.51 ± 0.10 pN. Treatment with pnBB or nocodazole reduced the tension to 1.43 ± 0.12 and 2.07 ± 0.15 pN respectively, while CK666 increased the tension to 4.01 ± 0.15 pN.

We validated these results with two engineered controls. A G67S mutation was introduced into Venus to generate the Dark Acceptor (DA) construct, disrupting Venus’s fluorophore and acting as a control without FRET and donor fluorescence only. The flexible linker connecting the FRET pair was replaced by five amino acids (5AA) in the 5AA control, forcing the FRET pair in close proximity and thus a high FRET state. Due to this design, both controls are expected to be tension insensitive (Fig. 3A). Indeed, in comparison to the WT module, which had a FRET efficiency of 20.44 ± 0.57%, DA engineered control had a decreased FRET efficiency of 1.35 ± 0.73%, while 5AA control had an increased FRET efficiency of 36.73 ± 0.82%. Treatment with para-nitroblebbistatin did not change FRET efficiency values (1.1 ± 0.78% for DA and 35.23 ± 0.81 for 5AA) (Fig. 3B).

**Figure 3.**
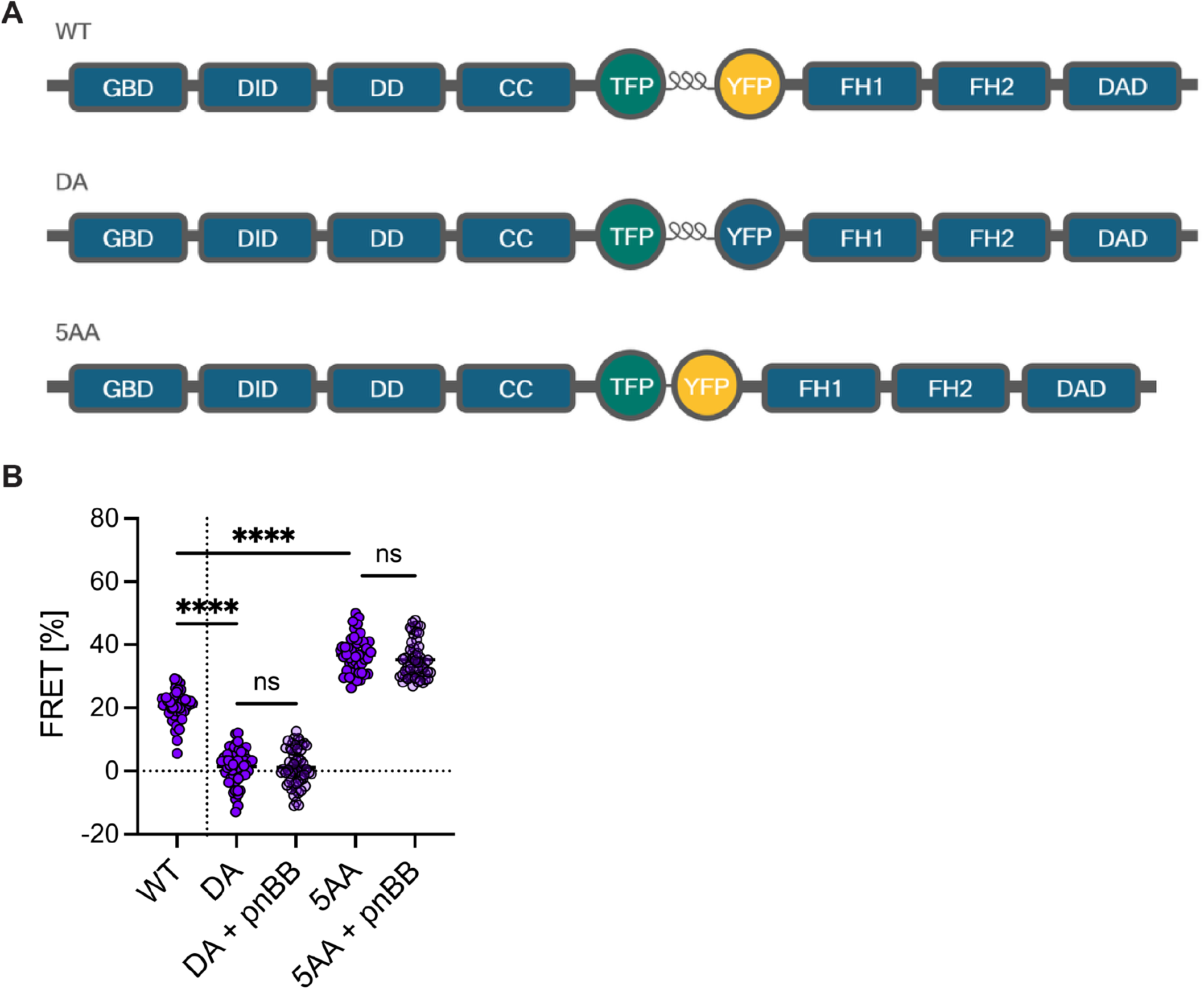
Design and characterization of FRET controls. **A –** Schematic of the WT mDia1 construct, the dark acceptor construct (DA: G76S mutation in YFP), and short linker construct (5AA: 5 amino acids replacing the flexible linker). **B –** Graph summarizing mean ± SEM of FRET percentage (% FRET) for each control in the presence or absence of para-nitroblebbistatin (pnBB). Each data point represents the FRET% obtained using whole-cell ROI. N = 50 – 65 cells per treatment, **** p <0.0001 based on ordinary one-way ANOVA with Tukey’s multiple comparisons test. Scale bar = 20 µm.

### M2 paralogs exert specific effects on tension transmitted towards mDia1TS

We next assessed the effects of non-muscle myosin 2 paralogs on tension. Since U2OS cells express all three paralogs of M2^36^, we used RNA interference to knock down each paralog individually (Fig. 4). While the raw fluorescence of mDia1TS was largely unaffected (Fig. 4A), the image fits of cells transfected with M2 paralog targeting siRNAs displayed reduced tension compared to the non-targeting control. To validate the efficiency of knockdowns, a Western Blot using antibodies specific for M2A, M2B and M2C confirmed the absence of the respective, endogenous protein bands (Fig. 4B, see also Fig. S2). We quantified the contribution to the total, exerted tension of each paralog by analyzing populations of whole-cell ROIs and estimated the tension change from 3.47 ± 0.13 pN for cells expressing scrambled siRNA control to 2.25 ± 0.17, 2.70 ± 0.13, and 2.58 ± 0.12 pN for cells transfected with M2A, M2B, and M2C siRNAs, respectively (Fig. 4C).

**Figure 4.**
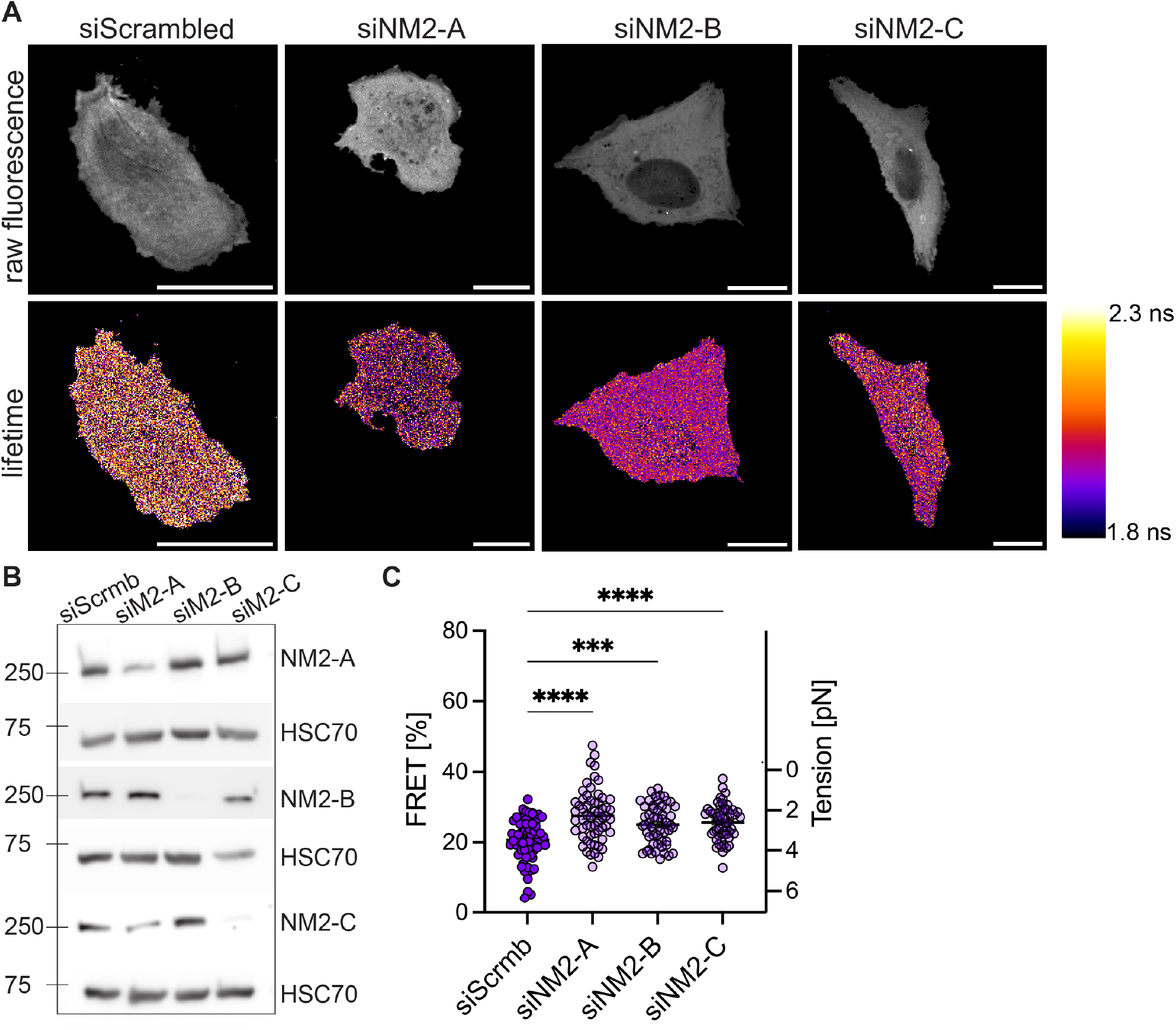
M2 paralogs exert specific effects on tension transmitted towards mDia1TS. **A –** Representative raw fluorescence (top) and lifetime color-coded (bottom) images of U2OS cells transfected with non-targeting scrambled siRNA or siRNAs against NM2 paralogs 2A, 2B, and 2C. **B –** An immunoblot from a representative experiment confirming knockdown of individual paralogs. **C –** Graph summarizing mean ± SEM of FRET percentage (% FRET) for each control siRNA- and M2 paralog siRNA-treated cells. FRET% was calculated using fits of whole-cell ROIs. The tension level of each paralog was calculated using TSMod’s calibration. N = 50 – 65 cells per treatment, *** p <0.001, **** p <0.0001 based on ordinary one-way ANOVA with Dunnett’s multiple comparisons test. Scale bar = 20 µm.

### Mitogen stimulation and physical constraints induced by micropatterning allow for measurement of redistribution of tension

To make the subtle variation in tension levels experimentally amenable within single cells and using mDia1TS, U2OS cells were examined on micropatterned coverslips. A crossbow-shaped area was functionalized with fibronectin, resulting in cell shapes resembling a growth cone with a distinguishable central and peripheral domain. The signal-to-noise ratio was improved by averaging over several patterned cells in the same conditions. The intensity profile of WT cells, plated in low serum conditions, showed a slight increase in the peripheral area compared to the central area, however, after acute cell stimulation with Epidermal Growth Factor (EGF), this difference became much more pronounced, indicating relaxation in the central area and tensing in the peripheral area. As a control to exclude optical artifacts, the tension sensor module (TSMod) alone was expressed, showing a flat lifetime profile throughout the entire cell (Fig. 5A-B). While these relative differences appear consistently, absolute values are difficult to compare due to the variation of lifetime levels from cell to cell; and are better achieved using whole-cell ROIs.

**Figure 5.**
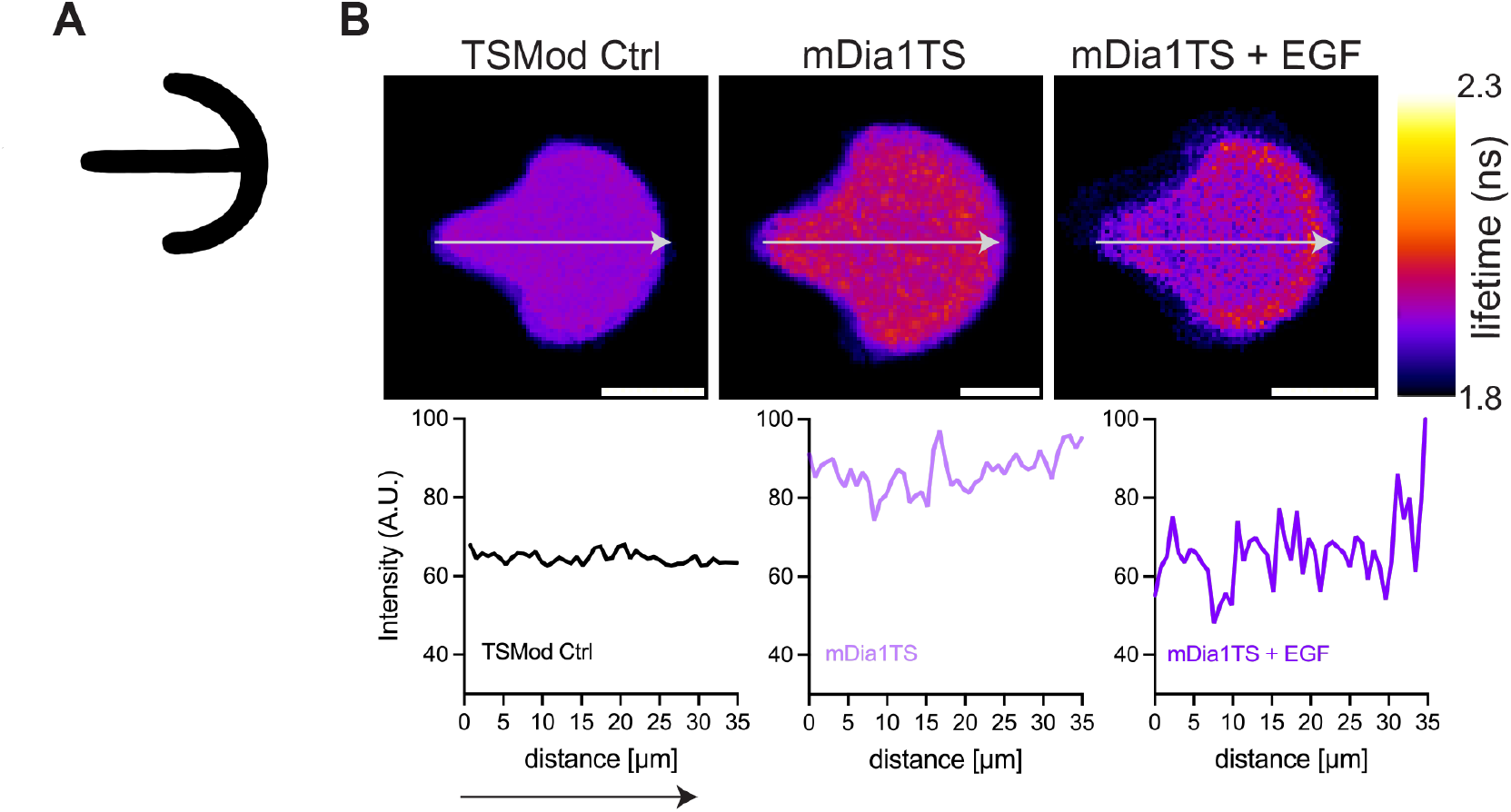
Micropatterned mapping reveals local mDia1TS FRET% fluctuations. **A –** Schematic of the 1100 µm^2^ crossbow pattern functionalized with fibronectin. **B –** Image fits of cells plated on micropatterns and intensity profiles along the cell averages. N = 7 – 10 cells per condition.

## Discussion

In this study, we designed, characterized and validated a FRET based tension sensor of mDia1, mDia1TS, and directly quantified reported tension in live U2OS cells. This probe offered direct evidence that mDia1 in cells is under load, which has been speculated given the tension-sensitive behavior that has been reported by several in vitro studies^20,37^. According to those studies, the polymerization rates of mDia1 accelerate when tension is applied via constrained actin filaments. Near maximum rates are reached before 2 pN, which is well before the baseline cortical tension reported by mDia1TS, which was ∼3.5 pN. Mechanistically, this behavior is explained by the ring-shaped mDia1 dimer alternating between “closed” and “open” conformations, resulting in a stair stepping-like mechanism of barbed-end polymerization. Tension presumably stabilizes the “open” confirmation, facilitating addition of G-actin monomers to the barbed end, and resulting in the accelerated dynamics of mDia1 under strain.

While the reported tension seems to be in the range of what has been reported from in vitro studies, a direct calibration of the sensor is missing for *bona fide* confirmation of absolute values. Here, a published calibration of the tension-sensitive linker was used, with corrections to adjust for differences in the dyes’ Förster radii (see Methods section). However, the data suggests that mDia1TS is excellently suited to measure relative values between populations of cells with pN precision. We leveraged mDia1TS to determine differences in the actin cortex’s tension after treating the cells with cytoskeletal drugs and by knocking down paralogs of M2, which allowed us to directly determine the contribution to the total tension from M2A, M2B and M2C respectively. The inhibitor of total M2, pnBB, was the cytoskeletal drug which led to the most dramatic reduction of the measured tension. It is noteworthy that even with pnBB, the tension was not reduced to zero, suggesting that the force balance mediated towards the actin cortex is complex and not mediated by M2 alone. This was further confirmed by the significant reduction of tension after addition of Nocodazole, a microtubule inhibitor. It has been reported previously that microtubules directly interact with mDia1, and the data reported here suggests mechanical feedback at the intersection with mDia1. We further validated these findings by using two tension-insensitive controls of mDia1TS, DA and 5AA, which simultaneously are reporters for the lower (zero FRET) and highest possible FRET values, by forcing the donor-acceptor pair in close proximity with a short, five amino acid linker. These controls were unaffected by the addition of pnBB, and confirmed that all measured FRET% using the WT construct were well between those two limits. This suggests that the sensor maxing out is unlikely under physiological conditions, albeit still possible.

Further limitations of the probe are set by the robustness of FLIM fitting, which in turn heavily depends on the photon count^38-40^. While the used U2OS cells appeared to be unaffected by higher expression level of the exogenous mDia1, low expression levels resulted in photon counts that were insufficient for double-exponential fitting. However, double-exponential fits were generally necessary for the decay curves from mDia1TS, as confirmed by χ2-tests. This meant that the signal was composed of two main components, and all lifetimes reported in this study were their intensity-weighted averages. This measure was chosen to reflect the biological meaning of these components, as the process of tensing and relaxation at the cortex, as well as the resulting FRET% are likely highly dynamic. We interpret the two lifetime components as the two extreme states of mDia1TS (“relaxed” and “tensed”, Fig. 1), meaning that the intensity-weighted average directly reflects the dynamic balance between these two states. It is expected that not all mDia1 molecules are engaged simultaneously, due to a population of inactive mDia1 that is not activated by RhoGTP and not mechanically anchored at the membrane. Evidence for this interpretation is found by the effect of CK666, which resulted in the only measured increase in FRET%. This can be explained by the inhibition of the Arp2/3 complex, freeing up actin filaments to engage with mDia1. This leads to a higher fraction of mDia1TS photons reporting the long (“tensed”) lifetime component, thus explaining the measured increase in FRET%.

While limited photon count and low signal-to-noise ratios generally masked the nuances of subcellular tension, we present two strategies that overcome these limitations. First, by maximizing the photon count using whole-cell ROIs and analyzing significant populations of cells under the same conditions. We used at least 40 cells per condition, as the lifetime variation from cell to cell was considerable after fitting the decay curves of these large ROIs. To determine if this variance was biologically relevant, we used a control, the tension sensor module alone (TSMod). The module alone, without any endogenous protein components reported comparable variance, which confirmed that this effect resulted from methodological noise rather than reflecting biological meaning, such as the inherent dynamics of tensing/relaxing. To show that subcellular, spatial nuances can be resolved using mDia1TS (Fig. 5), we used micropatterning of U2OS cells. This fixed the shape of the cells, which allowed us to average over several image fits under the same conditions, highlighting the trend of cells to report higher tension in their periphery and lower tension in the center. This spatial heterogeneity was remodeled after acute EGF stimulation, leading to further relaxation in the center, with a pronounced, tense periphery. It is noteworthy that analyzing whole-cell ROIs, the lifetime throughout the cell appeared unaffected, meaning that the average tension did not significantly change between WT and WT+EGF, even though the sub-cellular distribution showed increased polarity. This suggests that mDia1TS is sensitive enough to report tension spikes near the periphery, likely mediated by active RhoA/Cdc42 and periphery-localized focal adhesion complexes, and directly shows that mDia1 tracks this mechanical landscape in real time.

One aspect not addressed in this study is the effect of the position where TSMod is embedded. A position just in front of the actin-binding FH1/FH2 domains was chosen, where TSMod captures the immediate mechanical tension between the N-terminal membrane anchor and the unstructured, but extended FH1 domain. Due to the distance to functional domains, this position is unlikely to disturb dimer formation, actin binding or any of the autoregulatory functions of mDia1. However, data from fission yeast formin Cdc12 showed that the reported tension fluctuates depending on where the module is embedded^41^. While the chosen placement for mDia1TS is expected to be optimal to report the pulling forces within the actin cortex, it will be an interesting perspective to probe various insertion sites in future studies.

Taken together, this study finds mDia1TS as a non-invasive and site-specific tool to study mechanical tension in the actin cortex. Existing techniques, like atomic force microscopy (AFM), optical tweezers, or traction force microscopy measure bulk forces or extracellular traction forces^32,42-45^, while the data reported here reflects intracellular and molecule-specific tension experienced by mDia1. This opens the way for further studies, leveraging mDia1TS’s capabilities to report mechanical force fluctuations at the membrane-cortex intersection. We demonstrate that this is possible both comparing populations of cells and by averaging image fits to achieve subcellular localization. Potentially, this can be used for time-resolved studies such as during active cell migration or monitoring constriction events during endocytosis, or quantifying specific loci where mDia1 is abundant, such as in filopodia tips.

## Materials and Methods

### Chemicals and Reagents

Arp2/3 inhibitor CK666 (Selleckchem cat# S7690), microtubule depolymerization inhibitor nocodazole (Selleckchem cat# S2775), non-muscle myosin 2 inhibitor *para*-nitroblebbistatin (Cayman Chemicals cat# 24171) were prepared as 25 – 100 mM stock solutions in anhydrous DMSO (ThermoFisher cat# D12345) and stored at -80 ºC. Human EGF (PeproTech cat# AF-100-15-100UG) was prepared as a 100 µg/mL stock in 2% BSA in tissue-culture grade sterile PBS and stored at -20 ºC.

### Plasmids and Cloning

Plasmids containing mDia1 and the Tension Sensor module were acquired from addgene (plasmid #54157 and #26021). The plasmids containing controls (short linker: 5AA, dark acceptor: DA) were kindly provided by Dr. Indra Chandrasekar (Sanford Research). Overlapping fragments of mDia1 were PCR-amplified using forward primer 5’-CTAGCGTTTAAACTTAAGCTTATGCTAGTGACGTCAGATCCGC-3’ and reverse primer 5’-ACTCGCCCATCTTGTACAGCTCGTCCATGCC-3’ for the N-terminal fragment of mDia1, foward primer 5’-GCTGTACAAGATGGGCGAGTTTGACATCCG-3’ and reverse primer 5’-TGCTCACCATTTCGAGGGAACCTCTCTCGG-3’ for the tension sensor modules, and forward primer 5’-TTCCCTCGAAATGGTGAGCAAGGGCGAGG-3’ with reverse primer 5’-CCCTCTAGACTCGAGCGGCCGCCTAGATGCATGCTCGAGCGG-3’ for the C-terminal fragment of mDia1. The oligonucleotides were synthesized by Eurofins Genomics. Phusion DNA polymerase (New England Biolabs cat# M053) was used for all amplification reactions using the manufacturer’s recommended protocol, at an annealing temperature of 65°C and in the supplied Phusion GC buffer (New England Biolabs cat# B0519) supplemented with 3% DMSO and 2.5 M betaine. The amplified fragments were gel purified using a Monarch Spin DNA Gel Extraction Kit (New England Biolabs cat#T1120) and subsequently combined using In-Fusion seamless cloning (Takara Bio cat# 638947) into a linearized pcDNA3.1 mammalian expression vector and stored via transformation into Stellar Competent *E. coli* (Takara Bio cat# 636766) using the protocol included in the manufacturer’s manual. Sequences of resulting plasmids were verified using Plasmidsaurus, Inc.

### Cell lines and Transfection

U2OS cell line (ATCC) was cultured in DMEM, high glucose, GlutaMAX (Gibco cat# 10564011) supplemented with 10% heat inactivated FBS (Gibco cat# 16140071) and 1X antibiotic-antimycotic (Gibco cat# 15240062). Cells were transfected with the indicated plasmids using Lipofectamine 2000 (Invitrogen cat# 11668027) or JetPRIME (Polyplus cat# 101000015).

For siRNA-mediated M2 isoform-specific knockdown, U2OS cells were transfected with 50 nM non-targeting siRNA (D-001810-10-05) or siRNA against M2A (ON-TARGETplus Human MYH9 siRNA SMARTPool L-007668-00-0005), M2B (ON-TARGETplus Human MYH10 siRNA SMARTPool L-023017-00-0005), or M2C (ON-TARGETplus Human MYH14 siRNA SMARTPool L-027149-01-0005) using JetPRIME. Forty-eight hours post-siRNA transfection, cells were transfected with the indicated plasmids using Lipofectamine 2000.

### Immunoblotting

Efficiency of M2 isoform knockdown was evaluated by immunoblotting 72 hours post siRNA transfection. Cells were lysed in cell lysis buffer (CST cat# 9803) supplemented with a protease inhibitor cocktail (CST cat# 5871). Lysates were briefly sonicated and protein levels were quantified using BCA assay kit (Pierce cat# 23225). Lysates were resolved on 6% discontinuous Tris - Glycine SDS-PAGE gel and transferred to nitrocellulose membranes *via* Trans-Blot Turbo transfer system (Bio-Rad cat# 170415). Membrane slices were blocked in 3% milk in TBS, then incubated overnight in primary antibodies diluted in TBS + 0.05% Tween-20. The following dilutions were used: NM2-A (CST #49349, rabbit monoclonal), NM2-A (Sigma-Aldrich #M8064, rabbit polyclonal), NM2-B (Invitrogen #PA5-17026 rabbit polyclonal), NM2-B (BioLegend #909901, rabbit polyclonal), NM2-C (CST #3405, rabbit polyclonal), HSC70 (Santa Cruz Biotech #sc-7298, mouse monoclonal). The following day, the membrane slices were washed thrice in TBS-T and incubated in HRP-conjugated secondary antibodies (Jackson ImmunoResearch) for 1 h and room temperature. Membranes were processed on Amersham AI600 chemiluminescent imager and resulting image captures were further analyzed using Fiji/ImageJ.

### Cell Seeding and Treatments

Glass-bottom 35 mm dishes (Mat-Tek cat# P35G-1.5-14-C) were incubated with 20 µg/mL fibronectin (Sigma-Aldrich cat# F1141) in tissue-culture grade PBS at 4ºC overnight. Dishes were rinsed in PBS and equilibrated in media in a 37 ºC incubator prior to cell seeding. Cells were seeded at 20 – 30,000 cells per dish in imaging media (DMEM (no phenol red, high glucose, HEPES; ThermoFisher cat# 21063029) supplemented with 10% heat inactivated FBS (Gibco cat# 16140071), 1X antibiotic-antimycotic (Gibco cat# 15240062), and 50 µM ascorbic acid (Sigma-Aldrich A4544)).

Cells were allowed to spread for 4 hours prior to inhibitor treatment. Inhibitors or DMSO vehicle control dilutions were prepared immediately before use at a 2X concentration in imaging media. Final concentrations of inhibitors were as follows - CK666, 100 µM; nocodazole, 10 µM; pnBB, 25 µM.

For cell imaging on micropatterns, transfected cells were plated and serum-starved on fibronectin-coated crossbow-shaped medium micropattern coverslips (CYTOO, Inc.) in reduced-serum media ((DMEM (no phenol red, high glucose, HEPES; ThermoFisher cat# 21063029) supplemented with 2% heat inactivated FBS (Gibco cat# 16140071), 1X antibiotic-antimycotic (Gibco cat# 15240062), and 50 µM ascorbic acid (Sigma-Aldrich A4544)). Cells were allowed to spread on patterns for 4 hours prior to imaging. In experiments with EGF stimulation, cells were then incubated in reduced-serum media supplemented with EGF (50 - 100 ng/mL) for 30 min prior to image collection.

### FLIM-FRET Imaging

Fluorescence lifetime imaging (FLIM) was carried out using a Leica SP8 STED 3X/Confocal Microscope equipped with a pulsed white-light laser set to 470 nm and a Falcon FLIM system. Detection was performed with a Leica HyD SMD hybrid photomultiplier tube collecting quenched donor emission in the 480–520 nm range, along with a Leica HC PL APO CS2 63× 1.40 NA oil immersion objective (Leica Microsystems, Wetzlar, Germany). Images were acquired at a scanning speed of 400 Hz with a 512 × 512 pixel format and a pinhole diameter of 1.5 Airy Units. Before imaging, the laser power was adjusted to achieve fewer than 0.1 photons per laser pulse (approximately 5 µW at the back aperture). Frames were collected to achieve a photon count of at least 10^6^ per image.

### Analysis

FLIM images were analyzed in Leica Las X by manually drawing ROIs around single cells. Each ROI’s decay curve was fit using the n-Exponential Tail model in the range of 1 to 8 ns. All WT mDia1TS and 5AA mDia1TS measurements were fit with a double-exponential, while DA mDia1TS could be fit with a single-exponential. Higher exponentials did not improve the fits’ χ2 significantly. For all measurements, the readouts were the intensity-weighted, averaged lifetimes of its components, that were subsequently used to calculate FRET% using the formula

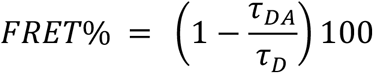

where τ_DA_ is the measured Donor lifetime in the presence of the acceptor, and τ_D_ is the average lifetime of the DA control (donor fluorescence only). To estimate the module’s tension, the FRET/force relationship of a published calibration curve was used, with a correction based on the Förster radius of the original Cy3/Cy5 FRET pair versus the fluorescent proteins (TFP/Venus) used in this study^31,46,47^. FLIM image fits were made using the same parameters in Leica Las X. For micropatterned cells, the binning was set to 4, then exported for further analysis in FIJI/ImageJ^48^. A stack of single-cell image fits was first registered using the StackReg plugin^49^, then averaged using an average intensity z-projection.

## Acknowledgements

The authors would like to thank the NHLBI Light Microscopy Core for training and support.

This research was supported by NHLBI DIR project number ZIAHL001786 to J.R.S., NHLBI DIR project number ZIAHL006121 to J.A.H. A.G. is supported by the NHLBI Lenfant Biomedical Fellowship and NLHBI K22HL179265 Career Transition Award.

The contributions of the NIH authors were made as part of their official duties as NIH federal employees, are in compliance with agency policy requirements, and are considered Works of the United States Government. However, the findings and conclusions presented in this paper are those of the authors and do not necessarily reflect the views of the NIH or the U.S. Department of Health and Human Services.

## Author contributions

Conceptualization – P.B. and A.G.; Methodology – P.B. and A.G.; Validation – P.B. and A.G.; Formal Analysis – P.B. and A.G.; Data Curation – P.B. and A.G.; Visualization – P.B. and A.G.; Supervision – J.R.S. and J.A.H.; Funding Acquisition – J.R.S. and J.A.H.

## Figure Legends

**Figure S1. Truncation of the GBD domain leads to mDia1TS aggregation. A –** Schematic of the WT mDia1TS construct and the ΔGBD mDia1TS construct. **B –** Examples of U2OS cells expressing ΔGBD mDia1TS. White arrows indicate aggregates. Raw fluorescence is shown.

**Figure S2, related to Figure 4. Uncropped chemiluminescent and white light images of membranes used to make Figure 4B**. Red rectangles indicate approximate location of the crop.

