## Supplementary figures and images for "Development and Characterization of a FRET-based Formin Tension Sensor in Living Cells"

### Supplemental Figure 1

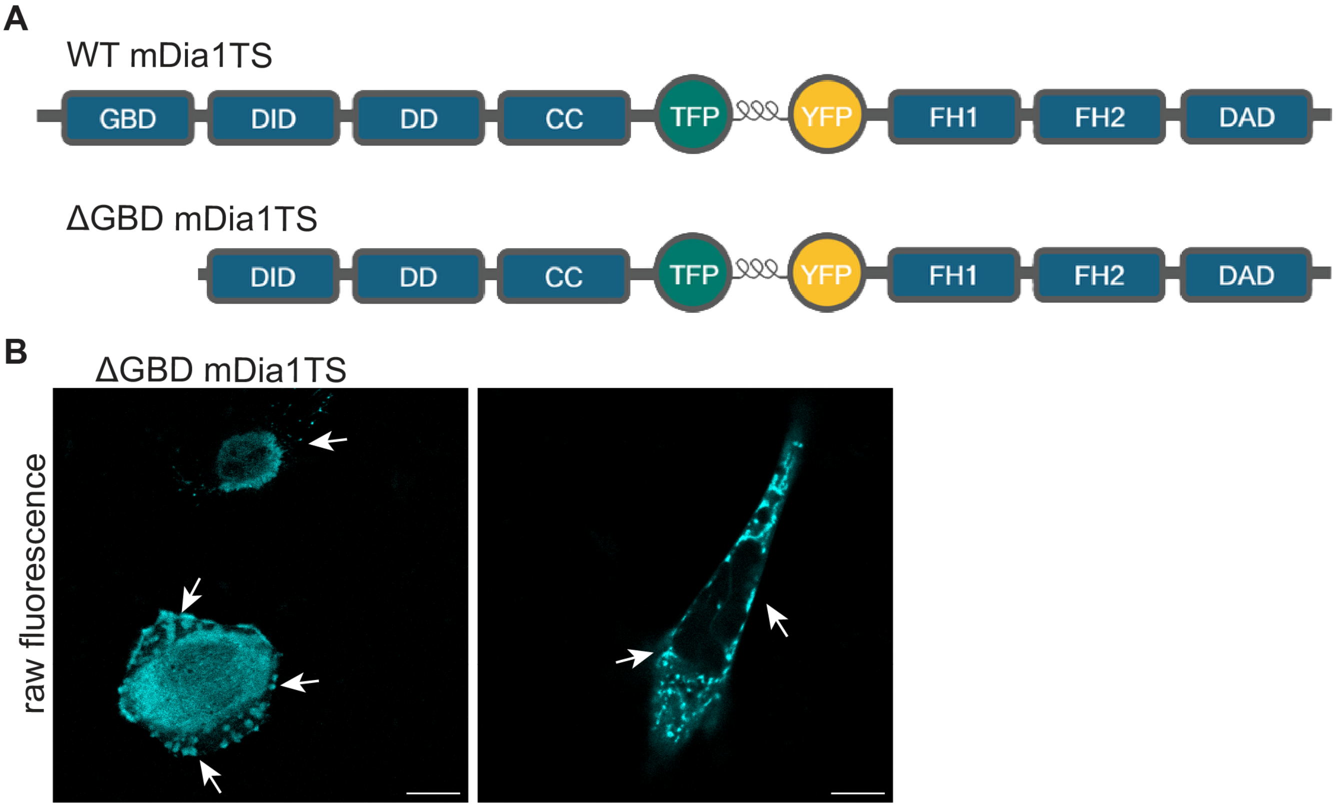

### Supplemental Figure 2

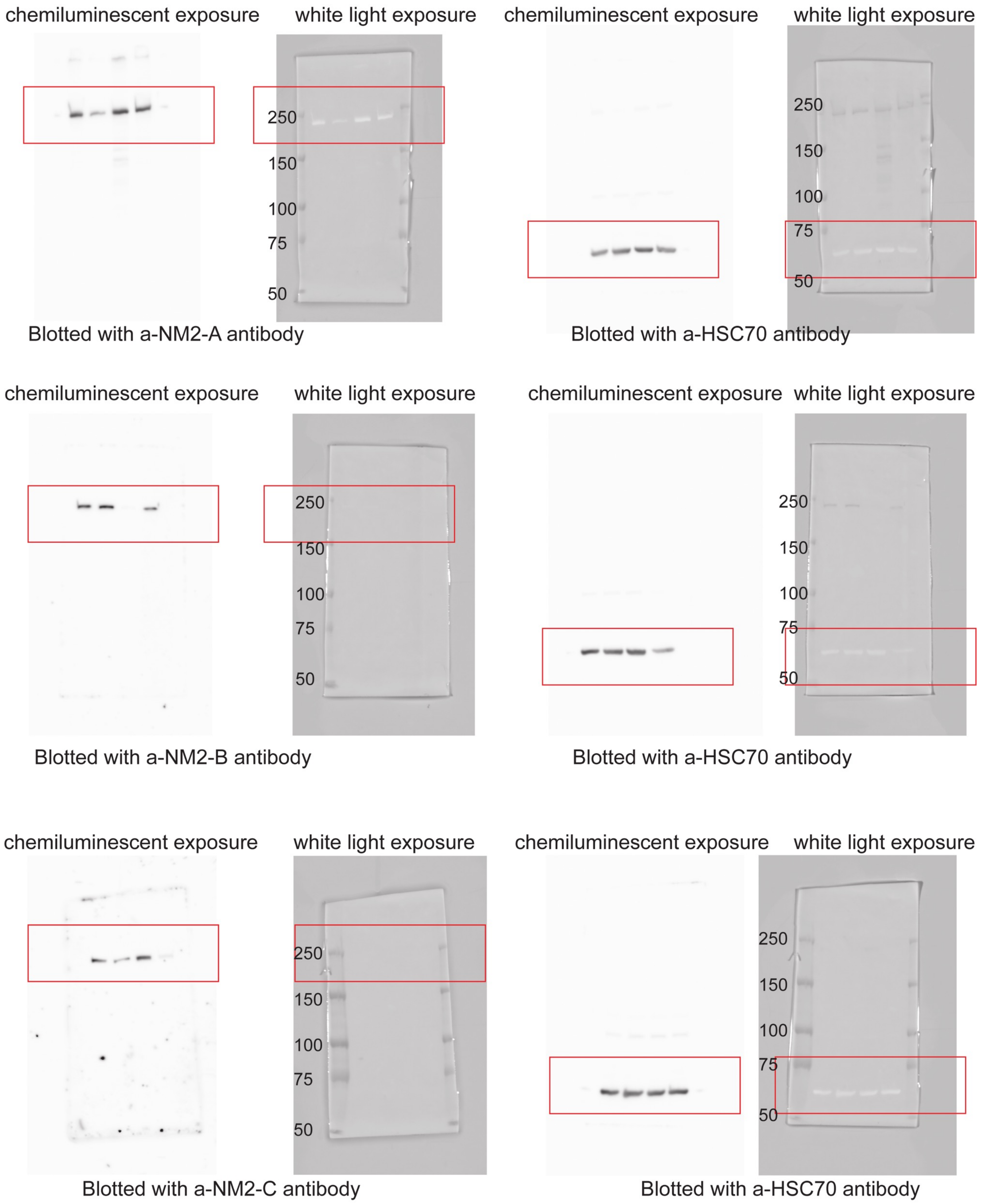
